# Increasing the time resolution of single-molecule experiments with Bayesian inference

**DOI:** 10.1101/099648

**Authors:** Colin D. Kinz-Thompson, Ruben L. Gonzalez

**Author notes:** To whom correspondence should be addressed (C.D.K.) and (R.L.G.).

## Abstract

Many time-resolved, single-molecule biophysics experiments seek to characterize the kinetics of biomolecular systems exhibiting dynamics that challenge the time resolution of the given technique. Here we present a general, computational approach to this problem that employs Bayesian inference to learn the underlying dynamics of such systems, even when they are much faster than the time resolution of the experimental technique being used. By accurately and precisely inferring rate constants, our Bayesian Inference for the Analysis of Sub-temporal-resolution Data (BIASD) approach effectively enables the experimenter to super-resolve the poorly resolved dynamics that are present in their data.

## Introduction

Given their inherent ability to eliminate ensemble averaging, time-resolved single-molecule biophysical methods have revolutionized the study of biological mechanisms by enabling distributions of molecular properties to be observed, stochastic fluctuations from equilibrium to be investigated, and transiently sampled reaction intermediates to be characterized (1). Generally, the majority of these methods involve making sequential measurements of an experimental signal that acts as a proxy for the underlying, time-dependent state of a biomolecule. As a result, this process yields a time-ordered series of discrete measurements from which the underlying dynamics of the corresponding biomolecule can be inferred (2). Unfortunately, the ability to resolve the continuously varying dynamics of the corresponding biomolecule from a series of discrete measurements is fundamentally limited. Indeed, whereas a biomolecule may exchange between multiple conformational states during a single measurement acquisition period, these states are collectively represented by a single, time-averaged measurement of the experimental signal. This effect is akin to chemical exchange effects in nuclear magnetic resonance (NMR) experiments, in which distinct resonance peaks can coalesce into a single, averaged resonance peak when a nucleus rapidly exchanges between distinct magnetic environments (3). As a result of this effect, many time-resolved, single-molecule biophysical methods often fail to detect or properly characterize mechanistically critical biomolecular processes that occur on or faster than the time resolution of the technique, including early steps in ligand binding, local folding events, and rapid conformational fluctuations (4, 5).

To push beyond the time-resolution limits of these single-molecule methods, we have developed a Bayesian inference-based computational approach, which we call Bayesian Inference for the Analysis of Subtemporal resolution Dynamics (BIASD), to infer the rate constants governing transitions between discrete states of a single molecule from the analysis of a time-resolved, single-molecule experimental signal-even if those rate constants are substantially faster than the time resolution of the recorded experimental signal. Much like learning the point spread function describing the fluorescence signal from a single fluorophore in a superresolution imaging experiment enables the spatial position of the fluorophore to be inferred beyond the spatial resolution of the experiment, learning the model describing the kinetic behavior of a single molecule in a time-resolved single-molecule experiment using BIASD enables the kinetic behavior of the single molecule to be inferred beyond the temporal resolution of the experiment. By using Bayesian inference, BIASD can also integrate information from other experiments to further enhance its resolving power, while also employing a natural framework with which to describe the precision that the amount of data collected during the singlemolecule experiment will lend to the determination of the parameters governing the single-molecule kinetics (2, 6, 7). It is worth noting that, in a close parallel to the approach we describe here, Bayesian inference has been previously employed to improve the time resolution of the time-dependent free induction decay in NMR spectroscopy experiments, resulting in an orders-of-magnitude improvement in spectral resolution (7, 8).

Here, we first describe the Bayesian inference-based framework underlying BIASD. We then use BIASD to analyze computer-simulated signal *versus* time trajectories (signal trajectories) and investigate the accuracy and precision with which we can infer the known rate constants for transitions between states that were used to generate the signal trajectories. We next use BIASD to analyze experimentally recorded fluorescence resonance energy transfer efficiency (E_FERT_) *versus* time trajectories (E_FERT_ trajectories) in order to infer the unknown rate constants for transitions between states in the E_FERT_ trajectories. Notably, the E_FERT_ trajectories that we have analyzed here had previously eluded analysis due to the presence of transitions that are much faster than the time resolution of the electron-multiplying charge-coupled device camera that was used to record them (9). Finally, we describe and demonstrate an extension of the BIASD framework that can be used to infer rate constants for experimental systems consisting of static or interconverting sub-populations of molecular properties within an individual or ensemble of molecules. Remarkably, we find that BIASD permits accurate inference of rates constants from time-resolved, single-molecule experiments, even when the rate constants are orders of magnitude larger than the time resolution of the signal trajectories.

## Bayesian Inference-based Framework Underlying BIASD

In biomolecular systems, functional motions—such as those involved in ligand binding and dissociation processes, or large-scale conformational rearrangements—very often involve the simultaneous formation and/or disruption of numerous, non-covalent interactions. The relatively low probability of simultaneously forming and/or disrupting these numerous interactions can therefore result in large, entropically dominated, transition-state energy barriers for such functional motions (10, 11). Consequently, individual biomolecules are generally expected to exhibit effectively discrete and instantaneous transitions between relatively long-lived states (5), an expectation that is consistent with the step-like transitions that are generally observed in time-resolved single-molecule experiments (12).

An important consideration when analyzing the signal trajectories from such single-molecule experiments is that whenever an individual molecule undergoes a transition from one state to another, the transition occurs stochastically during the time period, *τ*, over which the detector collects and integrates the signal to record a data point in the signal trajectory. Thus, the probability that a transition will coincide exactly with the beginning or end of the *τ* in which it takes place is essentially zero. As a result, when a transition takes place, the signal value that is recorded during that *τ* does not solely represent either of the states involved in that transition. Instead, it represents the average of the signal values corresponding to the states that are sampled during *τ*, weighted by the time spent in each of those states. This time averaging makes it imprudent to assign the signal value recorded during such a *τ* to any one particular state, a process called idealization, because the molecule will have occupied multiple states during that *τ*. Notably, when the rate constants for transitions between states become comparable to or greater than *τ*^−1^, there is a large probability that the *τ*s of a signal trajectory will contain one or more transitions, and that, consequently, many of the signal values of the signal trajectory will exhibit this time averaging. Given such a scenario, analysis methods in which individual *τ*s are assigned to particular states (*e.g.*, the widely used strategy of idealizing signal trajectories using signal thresholds (13), or hidden Markov models (HMMs) (14, 15)) will introduce significant errors into the calculated rate constants for transitions between states and into the signal values assigned to those states (2).

In order to overcome the potential errors associated with determining rate constants and signal values from the analysis of signal trajectories, BIASD instead analyzes a different parameter that depends upon the dynamics of the biomolecular system: the fraction of time that is spent in each state during the *τ*s in a signal trajectory (16–21). To illustrate this approach, consider the case of an individual molecule that undergoes stochastic, uncorrelated (*i.e.*, Markovian), and reversible transitions between two states, denoted 1 and 2, (*i.e.*, 1 ⇄ 2, with forward and reverse rate constants of *k*_1_ and *k*_1_, respectively) that have unique signal values of *∊*_1_ and *∊*_1_. If the fraction of time that the molecule spends in state 1 during a particular *τ* is *f*, then, because of the two-state nature of the system, the fraction of time that the molecule spends in state 2 during that *τ* is 1 − *f*. It is important to note that, although the molecule is in an equilibrium between states 1 and 2, the value of *f* for any particular *τ* will not necessarily be the equilibrium value of *f* = (1 + *k*_1_/*k*_2_)^−1^. This is because *τ* might not be long enough for sufficient time averaging to occur (*i.e.*, to invoke ergodicity). Instead, each *τ* will exhibit a distinct, time-averaged value of *f*.

The exact value of *f* for a particular *τ* will depend upon the molecule’s stochastic path through state-space during *τ*. As such, a probabilistic description of *f*, which accounts for all possible paths through state-space, is needed to describe the likelihood of observing a particular value of *f* during a *τ* (22). In particular, for the reversible, two-state system considered here, such a description, which has roots in the analysis of the NMR chemical exchange effects described above (23) and in sojourn-time probability distributions (24), was first given by Dobrushin (25). This particular expression (derived in the *Supporting Information*) is a function of *k*_1_, *k*_2_, and *τ*, and has been used in many single-molecule studies—though mostly in the context of photon counting experiments and without Bayesian inference-based implementations (16–26). Experimentally, if the exact values of *f*, *∊*_1_, and *∊*_2_ during each *τ* were known, one would be able to calculate the expected value of the corresponding time-averaged signal, *μ*, for each *τ*, because it would be the linear combination 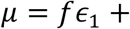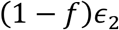. However, the analysis of time-resolved, single-molecule experiments deals with the opposite problem—observing a signal value, *d*, during each *τ* and trying to infer *f*, *∊*_1_, and *∊*_2_.

Generally, the values of *d* that are recorded during each *τ* are random variables, which are distributed according to a probability distribution function (PDF). For any number of states, this PDF for the observed values of *d* is the convolution of the PDFs for the signal values associated with each individual state, weighted by the fraction of time spent in that state (*i.e.*, *d*~*f*_1_*p*(*∊*_1_) * … * *f*_n_*p*(*∊*_*n*_), where *f*_*i*_ is the fractional occupancy of the i^th^ state, *p*(*∊*_*i*_) is the PDF of the signal values associated with the i^th^ state, and * denotes a convolution). For many experimental techniques, the signal values associated with each state are, or are approximately, distributed according to a normal PDF (*i.e.*, a Gaussian) with mean *∊*_*i*_ and variance 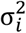 for the i^th^ state. Because the convolution of two normal PDFs is another normal PDF, in this case the PDF for the observed values of *d* is a normal distribution with mean μ = ∑_*i*_*f*_*i*_*∊*_*i*_ and variance 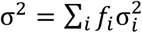. Furthermore, we can also account for noise from the detection process (*e.g.*, a normal PDF with mean 0 and variance 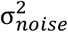), as well as a time-dependent baseline (*e.g.*, baseline drift at time *t*, *b*_*t*_, that is driven by white-noise is a normal PDF with mean b_*t*−1_ and variance 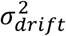 through additional convolutions; in these examples, the resulting PDF of *d* is again a normal PDF with mean μ = b_*t*−1_ + ∑_*i*_*f*_*i*_*∊*_*i*_ and variance 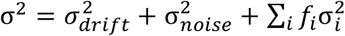. However, since *μ* and σ depend upon the set of fractional occupancies, {*f*}, which are not experimental observables, we have no way of knowing the exact form of this PDF—information that is required to calculate the probability of observing a particular value of *d*.

To circumvent this experimental limitation, the dependence of the PDF upon {*f*} can be removed by marginalizing {*f*} out of the expression for the PDF that was described above. This marginalized probability distribution of *d* then describes the likelihood of experimentally observing a particular value of *d* during a *τ* as a function of the set of rate constants for transitions between the states, {*k*}, the set of signal values corresponding to those states, {*∊*}, and the set of the amounts of noise in those states, {*σ*}, regardless of the exact values of {*f*} (Fig. S1A). As expected from the discussion in the previous section, this expression describes effects similar to those of chemical exchange in NMR experiments, in which rates with which nuclei exchange that are larger than the resonance frequency difference between exchanging nuclei cause distinct resonances to coalesce into a single, averaged resonance. As shown in Figure 1 for a two-state system (see *Supporting Information* Eqn. 10), the effect of increasing rate constants *k*_1_ and *k*_2_ results in distinct signal peaks centered at *∊*_1_ and *∊*_2_ to coalesce into a single, averaged peak centered at 〈*∊*〉.

**Figure 1:**
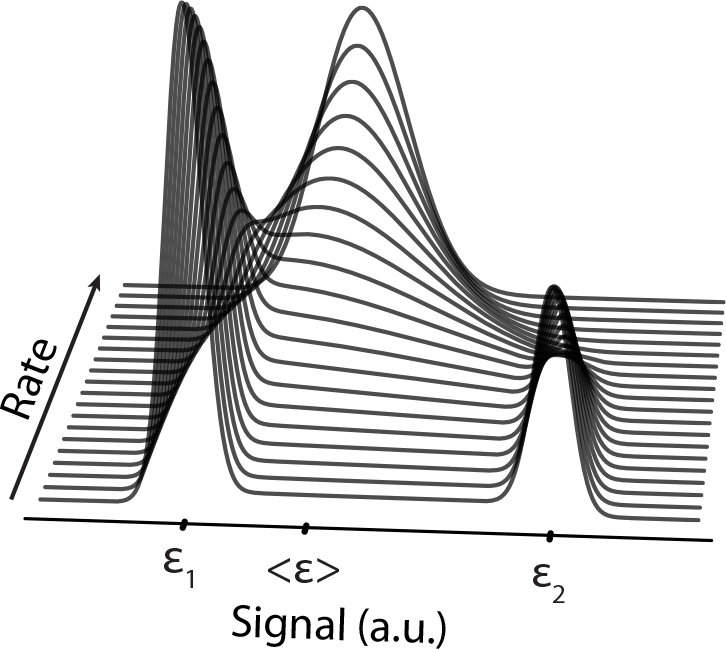
The marginalized, two-state likelihood function as a function of increasing rate of transitions. Arbitrary signal peaks at *ɛ*_1_ and *ɛ*_2_ coalesce into a single peak located at the equilibrium average 〈*ɛ*〉 as the rate constants for transitions between the two states increase relative to a fixed *τ*.

With such an expression describing the marginalized probability distribution of *d* we can then use Bayesian inference to estimate the parameters governing the single-molecule system (*i.e.*, {*∊*}, {*σ*}, and {*k*}) from the series of the *d* that comprise each of the signal trajectories. Primarily due to recent developments in computational tractability, Bayesian inference has become a powerful method for the analysis of biophysical data, such as determining the phases of X-ray reflections in X-ray crystallographic studies (27), performing simultaneous phylogenetic analysis of nucleotide and protein datasets (28), elucidating the number of structural classes present in cryogenic electron microscopy images (29), and ascertaining the number of states and the rates of transitions between those states present in single-molecule signal trajectories (30, 31). For an introduction to Bayesian inference, see Refs. 6, 7, 32, and the *Supporting Information*.

Unfortunately, performing Bayesian inference on a multi-parameter system, such as the one described here, results in a multi-dimensional, joint-probability distribution of the model parameters known as a posterior probability distribution, which is difficult to evaluate (32). In order to overcome this difficulty, we have chosen to evaluate the posterior probability distribution of the model parameters by numerically sampling it using a Markov chain Monte Carlo (MCMC) (6, 33) method with affine-invariant ensemble sampling (34, 35). Although alternative methods that approximate the posterior probability distribution of the model parameters, such as the Laplace approximation or variational inference, might be more computationally tractable, MCMC sampling is advantageous in that, unlike such approximation methods, it can provide an exact result that does not assume a particular structure of the posterior probability distribution (6). Regardless of the choice of method, however, the most important aspect of the approach described here is that we can evaluate the posterior probability distribution of the model parameters from the series of *d* that comprise a single-molecule signal trajectory in a manner that analytically accounts for the time resolution of the experimental technique.

In order to quantify the performance of BIASD and maximize its usefulness to the experimentalist community, we have also conducted a comprehensive analysis of how the posterior probability distribution behaves as a function of the parameters of the input signal trajectories (*c.f.*, *Dependence of BIASD Performance on Parameter Values in Supporting Information*). This analysis reveals that collecting additional data points in a signal trajectory increases the performance of BIASD, as does optimizing the sensitivity of the instrumentation so as to increase the signal-to-noise ratio (SNR) of the signal trajectory. While the results of these analyses can be used to determine the signal trajectory lengths and/or SNRs that would be needed to accurately and precisely infer rate constants that are some particular factor greater than *τ*^−1^, we find that the lengths and SNRs of the signal trajectories obtained using typical single-molecule instrumentation render BIASD useful for characterizing dynamics governed by rate constants that are up to three orders of magnitude greater than *τ*^−1^.

## Results and Discussion

### Analysis of Computer-Simulated Single-Molecule Signal Trajectories Reporting on the Kinetics of a Ligand Binding and Dissociation Process

To demonstrate the use of the analytical formulas underlying BIASD, we analyzed simulated singlemolecule signal trajectories that mimic the binding and dissociation of a ligand to its target biomolecule, a receptor, using the two-state, reversible kinetic scheme discussed in the previous section (36). In this example, ∊_1_ and ∊_2_ represent the signal values of the receptor in the ligand-free state and the ligand-bound state, respectively, and σ represents the standard deviation of the signal values for both states. Correspondingly, *k*_1_ and *k*_2_ represent the pseudo-first-order rate constant of ligand binding to the receptor, and the first-order rate constant of ligand dissociation from the receptor, respectively. As such, *k*_1_ is dependent on [*L*] with a dependence that is given by 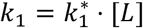, where 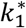 is the second-order rate constant for binding of the ligand to the receptor and [*L*] is the ligand concentration, and *k*_2_ is not dependent on [*L*]. In order to emulate a titration experiment, we varied the [*L*] to alter the fraction of ligand-bound receptor from ~0.1% to ~99.9% and simulated a series of individual signal trajectories where the [*L*] spanned six decades centered around the [*L*] corresponding to the equilibrium dissociation constant, [*L*] = *K*_*D*_. Notably, as is always the case for experimentally recorded signal trajectories, the finite length of each simulated signal trajectory presents an intrinsic limit to the amount of kinetic information contained in each signal trajectory. Finally, estimates of the parameters that were used to simulate the signal trajectories were obtained by analyzing the simulated signal trajectories using two idealization-based approaches: (i) half-amplitude signal thresholding (13) followed by Bayesian inference to infer the transition probability and quantify the kinetics (2) (Bayesian threshold), and (ii) an HMM (6) that used the maximum-likelihood framework to estimate the transition probability and quantify the kinetics (ML HMM). In addition to these two idealization-based approaches, the simulated signal trajectories were also analyzed using the BIASD approach presented here.

As shown in Figure 2A, the values of *k*_1_ and *k*_2_ obtained using both idealization approaches are inaccurate (green and red curves, respectively). Interestingly, however, neither approach absolutely outperforms the other in accuracy, and both plateau at the acquisition rate (*i.e.*, *τ*^−1^ = 10 s^−1^). Notably, the use of Bayesian inference in the Bayesian threshold approach tempers the fluctuations that are seen in the rate constants obtained from the ML HMM at high [*L*]. These fluctuations originate from the maximum likelihood framework used to estimate the transition probability in the ML HMM approach. Tempering of these fluctuations in the Bayesian threshold approach results from the use of a prior probability distribution, which describes the initial knowledge of the model parameters, in Bayesian inference (2, 6). Regardless, the rate constants were systematically underestimated across nearly the entire range of [*L*]s that were simulated, and this underestimation worsens with increasing [*L*]. It is striking that the values of *k*_1_ and *k*_2_ obtained using the Bayesian threshold idealization are also relatively precise—a misleading consequence of using idealization methods in general (2).

**Figure 2:**
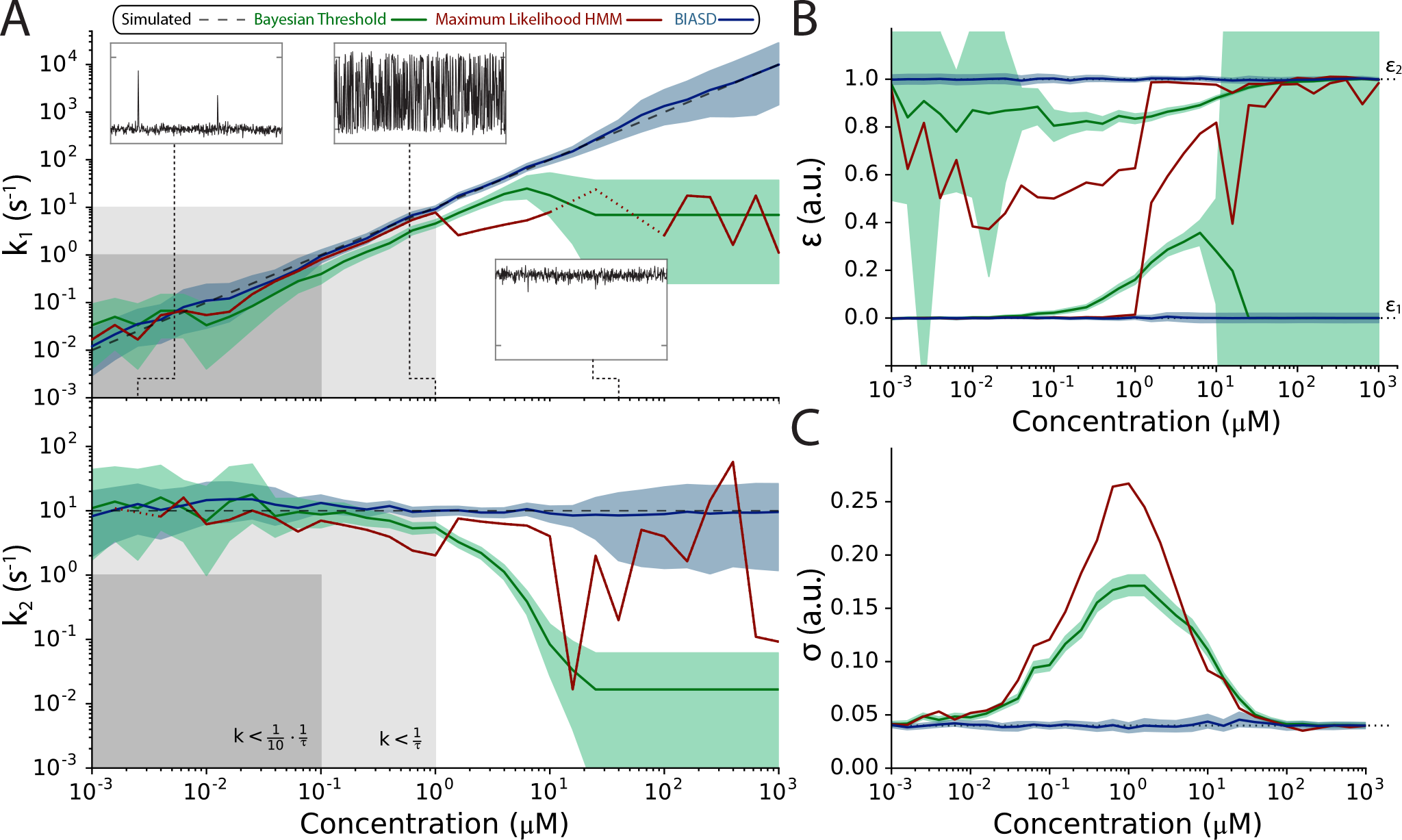
Analysis of *k*_1_ and *k*_2_ using BIASD (blue) and idealization-based (green and red) methods for a computer-simulated titration of a ligand to a receptor. [L] was varied three decades above and below the concentration where the equilibrium occupation probability of both states is equal to 0.5 (*i.e.*, K_D_ = 1). (A) Analysis of estimated rate constants *k*_1_, and *k*_1_. The regions where the rate constants are less than 1/10^th^ of the acquisition rate, *τ*^−1^, is shown in dark grey; the regions where the rate constants are less than the acquisition rate are shown in light grey. The simulated rate constants are plotted as the black dashed lines. The red line denotes the maximum-likelihood HMM estimate of the rate constants; dotted red lines indicate interpolated values due to transition probability estimates of unity. The green and blue areas denote the 95% credible intervals of the posterior probability distributions from analysis with half-amplitude thresholding-based Bayesian transition probability analysis (2, 13), and BIASD, respectively. Insets show the simulated signal trajectory corresponding to the indicated concentration. (B) Analysis of estimated signal values *ɛ*_1_, and *ɛ*_2_. Simulated values are plotted as black dashed lines. (C) Analysis of estimated signal noise *σ*

With regard to the values of ∊_1_ and ∊_2_ obtained using both idealization approaches, Figure 2B demonstrates that, while these methods can accurately determine the value of ∊_1_ if the receptor preferentially occupies the ligand-free state (low [*L*]) or ∊_2_ if the receptor preferentially occupies the ligand-bound state (high [*L*]), the time averaging caused by large values of *k*_1_ shift the inferred values of ∊_1_, sometimes quite significantly, toward the simulated value of ∊_2_, and *vice versa*. Here, the Bayesian threshold approach provided more accurate estimates of ∊_1_ and ∊_2_ than did the ML HMM. However, this was only because the signal threshold was set halfway between ∊_1_ and ∊_2_ using the known simulation parameters, thereby optimally minimizing the misclassification of states 1 and 2, and, consequently, maximizing the accuracy with which ∊_1_ and ∊_2_ are estimated; this is also why the estimates of σ from the Bayesian threshold approach are more accurate than those from the ML HMM (Fig. 2C) (13).

In contrast to both idealization approaches, the values of *k*_1_ and *k*_2_ obtained using BIASD are highly accurate (Fig. 2A). The simulated values of *k*_1_ and *k*_2_ are well encompassed by the 95% credible interval of the posterior probability distribution across the entire range of [*L*]s that were simulated, which includes rate constants that are up to three orders of magnitude larger than the simulated time resolution. In addition, the values of *k*_1_ and *k*_1_ are highly precise, as the 95% credible intervals of the posterior probability distribution are strikingly narrow over a range of [*L*] s corresponding to a value of *k*_1_ that is over an order of magnitude smaller than *τ*^−1^ to one that is over an order of magnitude larger than *τ*^−1^. Importantly, as the amount of data that is analyzed increases, the contribution that the choice of prior probability distribution (*i.e.*, the initial knowledge of *k*_1_, *k*_2_, ∊_1_, ∊_2_, and *σ*) makes to the posterior probability distribution diminishes. Consistent with the amount of data that are typically analyzed in single-molecule biophysical experiments, the results reported here are relatively insensitive to the prior probability distributions used for the analysis.

At the lower [*L*]s, the broadening of the posterior probability distribution that limits the precision for the estimates of *k*_1_ and *k*_2_ in both BIASD as well as the Bayesian threshold idealization arises from the finite amount of information regarding *k*_2_ and ∊_2_ that is contained in signal trajectories that exhibit very low occupation of the ligand-bound state of the receptor. Likewise, at the higher [*L*]s, the broadening of the posterior probability distribution, and the implied limitations to the precision for estimating *k*_1_ and *k*_2_ that is observed, arises from the finite amount of information regarding *k*_1_ and ∊_1_ that is contained in signal trajectories that exhibit very low occupation of the ligand-free state of the receptor. However, this broadening is somewhat attenuated, because the posterior probability distribution maintains consistency with the amount of previously known information about the underlying system contained within the prior probability distribution (*c.f.*, *Analysis Using BIASD in Supporting Information*). Regardless, the uncertainty at the lower and higher [*L*]s is a consequence of the finite amount of information in a finite-length signal trajectory, as many reciprocal pairs of *k*_1_ and ∊_1_ values (*i.e.*, a larger ∊_1_ and a smaller ∊_1_ or a smaller *k*_1_ and a larger ∊_1_) are consistent with the data. In an experimental situation, this imprecision can be alleviated by employing prior probability distributions for the {∊} values using the results of experiments performed under conditions in which one state is preferentially occupied, for instance, the values of ∊ observed in the absence of ligand could be used to construct a prior probability distribution for the values of ϵ associated with the ligand-free state of the receptor, whereas the values of ∊ observed in the presence of saturating [*L*] could be used to construct a prior probability distribution for the values of ∊ associated with the ligand-free state of the receptor. In the case of large-scale conformational rearrangements, one could similarly use a buffer condition, ligand, temperature, or mutation that preferentially stabilizes one state, or, alternatively, one could use molecular structures or models to estimate prior probability distributions of {∊} values. With regard to the values of ∊_1_ and ∊_2_ obtained using BIASD, Figure 2B demonstrates that these values were accurately inferred regardless of the value of [*L*], even at [*L*]s at which the idealization approaches drastically misestimate them. Finally, unlike the idealization approaches, which were only able to successfully infer σ when the signal trajectories were almost entirely in the ligand-bound-or ligand-free states, BIASD was also able to accurately and precisely infer σ from the simulated signal trajectories with intermediate values of [*L*] (Fig. 2C).

In summary, we were able to use BIASD to obtain accurate and precise posterior probability distributions for *k*_1_, *k*_2_, ∊_1_, ∊_2_, and *σ* across the entire range of [*L*]s that were simulated. Notably, BIASD was even successful when the rate constants in the simulated, single-molecule signal trajectories were much smaller then *τ*^−1^ although we note that, in this regime, the conventional analysis of idealizing the signal trajectories is much more computationally efficient. Most importantly, BIASD was able to accurately and precisely infer the rates constants and the signal values for simulated, single-molecule signal trajectories in which the rate constants were up to three orders of magnitude larger than *τ*^−1^ and up to about four orders of magnitude larger than the *τ*^−1^s where conventional idealization of signal trajectories begins to yield significant errors in the rate constants. This finding is consistent with the expected performance of BIASD (*c.f.*, *Dependence of BIASD Performance on Parameter Values in Supporting Information*).

### Analysis of Experimentally Observed Single-Molecule E_FRET_ Trajectories Reporting on the Kinetics of a Large-Scale Conformational Rearrangement

To demonstrate the use of BIASD in the analysis of experimental data, we chose to analyze experimentally observed, single-molecule E_FRET_ trajectories reporting on a large-scale conformational rearrangement of the ribosome. This essential, two-subunit, ribonucleoprotein-based, biomolecular machine is universally responsible for the translation of messenger RNAs (mRNAs) into proteins in living cells. The ribosome synthesizes proteins by repeatedly incorporating amino acids, delivered in the form of aminoacyl-transfer RNA (tRNA) substrates, into a nascent polypeptide chain in the order dictated by the mRNA being translated. During the elongation stage of protein synthesis (37), the ribosomal pre-translocation (PRE) complex undergoes stochastic, thermally driven fluctuations between two major, on-pathway conformational states that we refer to as global state 1 (GS1) and global state 2 (GS2), defining a dynamic equilibrium, GS1⇌GS2 (38, 39). These transitions between GS1 and GS2 constitute large-scale rearrangements of the PRE complex that involve relative rotations of the ribosomal subunits, reconfigurations of the ribosome-bound tRNAs, and repositioning of a ribosomal structural domain known as the L1 stalk (Fig. 3A) (40).

**Figure 3:**
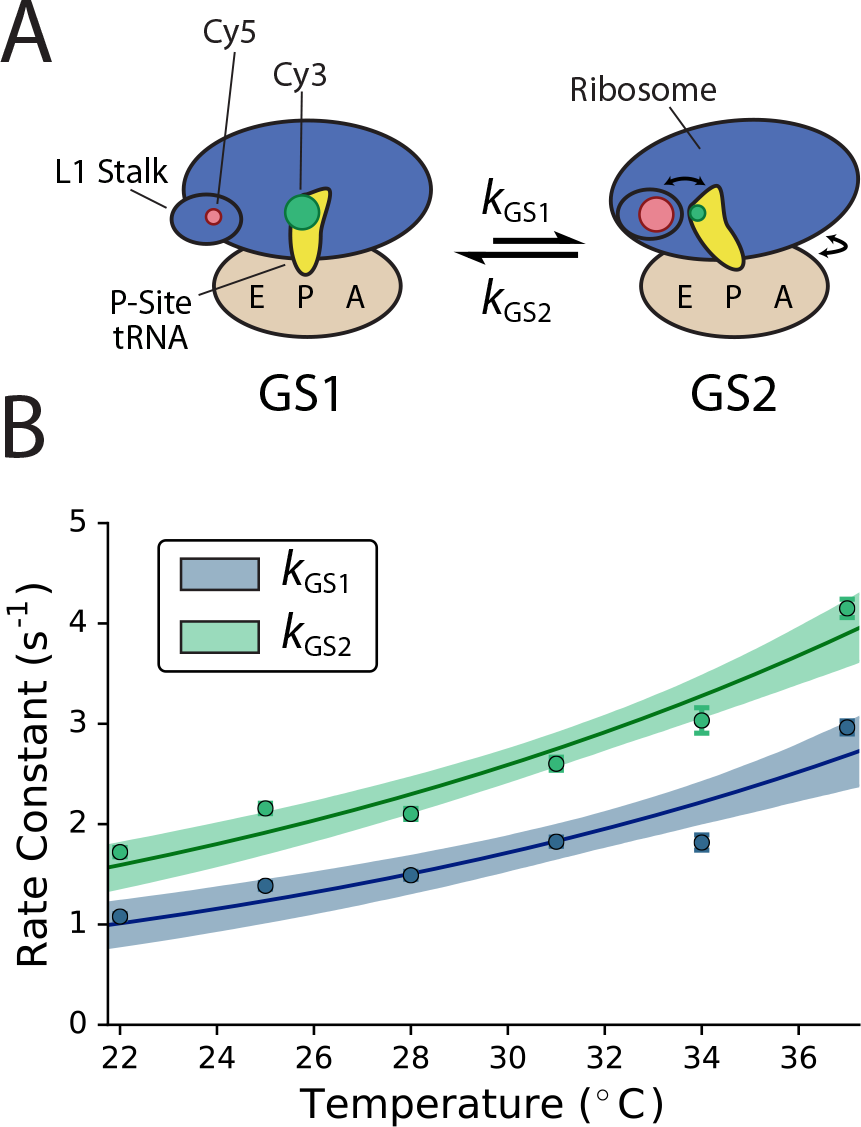
(A) Cartoon schematic of the GS1 ⇌ GS2 equilibrium on the PRE^−A^ complex previously studied by Wang and coworkers (9). Approximate positions of the Cy3 FRET donor and Cy5 FRET acceptor fluorophores of the ‘L1-tRNA’ labeling scheme used by Wang and coworkers are shown as green and red circles, respectively. The size of the fluorophores denotes the relative fluorescence intensity of each fluorophore in each state due to FRET. A, P, and E denote the A, P and E sites of the ribosome, respectively. (B) Temperature dependence of *k*_GS1_ and *k*_GS2_ for PRE^−A^ complexes using BIASD. The scatter plots show the expectation value of the posterior probability distributions of *k*_GS1_ and *k*_GS2_ and the error bars represent the 95% credible interval. The solid lines denote expectation values, and the shaded regions denote the 95% credible interval of the predictive posterior probability distribution from the transition-state theory analysis.

Previously, we have conducted wide-field microscopy smFRET studies of the temperature dependence of the rate constants governing GS1→GS2 and GS2→GS1 transitions by imaging a Cy3 FRET donor fluorophore-and Cy5 FRET acceptor fluorophore-labeled PRE complex analog lacking a tRNA at the ribosomal aminoacyl-tRNA binding (A) site (PRE^−A^) in a temperature-controlled, microfluidic observation flowcell (9). In these experiments, the increase in thermal energy that accompanied the increasing temperature caused the rate constants for the transitions between GS1 and GS2 to increase such that, at the highest temperatures, the E_FRET_ trajectories contained a significant number of time-averaged data points at the *τ* = 50 msec time resolution of the experiment (Fig. S2). Regrettably, the time averaging in these E_FRET_ trajectories precluded accurate determination of the rate constants, and, correspondingly, an analysis of the thermodynamic properties of the transition-state energy barriers that control the GS1→GS2 and GS2→GS1 conformational rearrangements (9). To overcome these limitations, we have used BIASD to analyze the sets of E_FRET_ trajectories of PRE^−A^ complexes that we have previously collected at 22, 25, 28, 31, 34, and 37 °C (9).

Here, we assume that the GS1⇌GS2 equilibrium can be represented by a single, reversible two-state kinetic scheme (Fig. 3A). In this kinetic scheme, *k*_*GS*1_ and *k*_*GS*2_ represent the unimolecular rate constants for the GS1→GS2 and GS2→GS1 conformational rearrangements, respectively. Correspondingly, ∊_*GS*1_ and ∊_GS2_ represent the E_FRET_ values of GS1 and GS2, respectively.

These six sets of E_FRET_ trajectories were analyzed using BIASD to provide estimates of *k*_*GS*1_, *k*_*GS*2_, ∊_GS1_, ∊_GS2_, and σ that best describe the entire set of E_FRET_ trajectories observed at each temperature. The values of *k*_*GS*1_ and *k*_*GS*2_ that were inferred using BIASD increase with temperature (Fig. 3B), and, at the highest temperatures, were greater than 1/10^th^ of *τ*^−1^ (*i.e.*, 2 s^−1^)-the regime where idealization approaches begin to systematically underestimate rate constants. We note that although the values of *k*_*GS*1_ and *k*_*GS*2_ inferred using BIASD are those that best describe the entire set of E_FRET_ trajectories observed at a particular temperature, inspection of individual E_FRET_ trajectories suggests the presence of kinetic heterogeneity as some are consistent with rate constants > 45 s^−1^, while others are consistent with rate constants < 0.1 s^−1^. This broad range of kinetic behaviors suggests the possibility that the PRE^−A^ complexes are compositionally heterogeneous (*e.g.*, sub-populations of PRE^−A^ complexes that differ in the aminoacylation status of the tRNA at the ribosomal peptidyl-tRNA binding (P) site, the presence or absence of a tRNA at the ribosomal tRNA exit (E) site, and/or the presence or absence of a particular ribosomal protein) and/or are conformationally heterogeneous due to structural rearrangements that are slow on the time-scale of the experiment and effect *k*_*GS*1_ and *k*_*GS*2_, but not necessarily ∊_GS1_ and ∊_GS2_. Additionally, we note that the posterior probability distributions of ∊_GS1_ and ∊_GS2_ that were inferred using BIASD have means of 0.13 and 0.78, respectively, which are values of ∊_GS1_ and ∊_GS2_ that very closely match the values of the mean E_FRET_ of GS1 and GS2 reported in previous, room-temperature studies of the analogous PRE^−A^ complex (0.16 and 0.76, respectively) (41). This correspondence strongly suggests that the values of ∊_GS1_ and ∊_GS2_ inferred using BIASD are accurate, regardless of the presence of time averaging in the *E*_FRET_ trajectories recorded at the highest temperatures.

With the inferred values of *k*_GS1_ and *k*_GS2_ as a function of temperature, we then used transition-state theory to quantify the apparent transition-state energy barriers along the apparent GS1→GS2 and GS2→GS1 reaction coordinates (42–45). Kramers’ barrier-crossing theory, which was developed to analyze thermally activated, condensed-phase transitions of a Brownian particle (43–45) and is increasingly being used to analyze the conformational dynamics and folding of small, globular proteins (12, 46), may ultimately provide a more exact analysis of the apparent transition-state energy barriers along the apparent GS1→GS2 and GS2→GS1 reaction coordinates. However, its application requires knowledge regarding the viscosity of the aqueous buffer in which the PRE^−A^ complex is dissolved and the ‘internal friction’ of the PRE^−A^ complex that are unavailable in the current study (12, 47). As such, we have opted to use transition-state theory, and regard the results as upper limits on the apparent transition-state energy barriers along the apparent GS1→GS2 and GS2→GS1 reaction coordinates, which do not account for internal friction or transition-state recrossings. To apply transition-state theory, we used the marginalized posterior probability distributions of the rate constants at each temperature to infer Δ*H*^‡^ and Δ*S*^‡^ from the equation *k*_*TST*_ = *kK*_*B*_*T*/*h exp*(−(Δ*H*^‡^ − *T*Δ*S*^‡^)/(*k*_*B*_*T*)), where *κ* is the transmission coefficient and is taken to be unity, *k*_*B*_ is the Boltzman constant, *h* is Planck’s constant, and Δ*H*^‡^ and Δ*S*^‡^ are the enthalpic and entropic differences between the transition and ground states (Fig. 3B). The marginalized results for the GS1→GS2 transition are 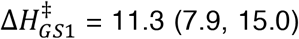 kcal mol^−1^ and 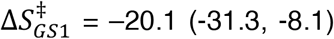 cal mol^−1^ K^−1^, and for the GS2→GS1 transition are 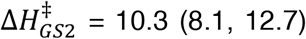 kcal mol^−1^ and 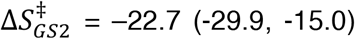 cal mol^−1^ K^−1^, where the numbers in parenthesis represent the lower and upper bounds for the 95% credible interval. Notably, the posterior probability distributions for Δ*H*^‡^ and Δ*S*^‡^ are highly correlated such that across all of the temperatures measured here, 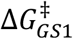 and 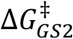 were sufficiently resolved; for instance, at 37 °C, 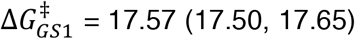 kcal mol^−1^, and 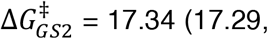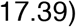 kcal mol^−1^. Structure-based interpretation of the absolute Δ*H*^‡^ and Δ*S*^‡^ values for the GS1→GS2 and GS2→GS1 transitions of a single PRE^−A^ complex is significantly complicated by the complexity of the enthalpic and entropic changes that are associated with conformational rearrangements of large macromolecular complexes, and by the inherent limitations of transition-state theory (5, 42, 48). Nonetheless, structure-based interpretations of the relative changes of the Δ*H*^‡^s and Δ*S*^‡^s (ΔΔ*H*^‡^s and ΔΔ*S*^‡^s) between different pairs of PRE^−A^ complexes (*e.g.*, containing different tRNAs at the peptidyl-tRNA binding (P) site, containing wildtype or mutant P-site tRNAs, consisting of wildtype or mutant ribosomes, *etc.*) are much more straightforward and can reveal the thermodynamic contributions that particular structural features of tRNAs or ribosomes make to the apparent transition-state energy barriers along the apparent GS1→GS2 and GS2→GS1 reaction coordinates. Combined with the temperature-controlled, single-molecule microscopy platform that we have previously described (9), the analytical framework presented in this section now enables the collection, analysis, and interpretation of such data.

### Inferring rate constants and signal values from systems with sub-populations of molecular properties

BIASD can be extended to address the presence of multiple, time-averaged sub-populations of molecular properties. These sub-populations may be static or interconvert, and may be present in an individual molecule or found among an ensemble of molecules. In such a situation, we can classify each data point as belonging to one of K different types of time-averaged sub-populations, and then use a ‘1-of-K’ vector, 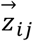, to denote to which of the K sub-populations the i^th^ data point from the j^th^ molecule belongs. Given the one particular sub-population specified by 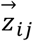, the likelihood of this data point being described by the parameters of this sub-population is calculated as described above for the case of the time-averaged, single-population system. Unfortunately, in an experimental situation there is no way of knowing which sub-population a particular data point belongs to, thereby preventing the likelihood of this data point from being evaluated; this situation is similar to that of the unknown fractional occupancy, *f*, described above.

To address this shortcoming, we could try to infer the values of all the 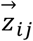 along with all of the other BIASD model parameters, but this is an unreasonable number of variables for an inference procedure. Additionally, we are often not concerned with the exact values of 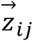, so much as with the occupancies of the K states (*e.g.*, the steady-state occupation probabilities) or with the rate constants that describe transitions between the K states. Fortunately, instead of performing inference to learn the model parameters and the set of 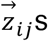, 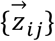, we can marginalize out all of the 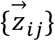 with the expressions for the probability of each 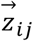. For instance, in the case of a mixture of static sub-populations of molecular properties among an ensemble of molecules (*e.g.*, a mixture of post-transcriptionally or post-translationally modified and unmodified molecules within an ensemble), these probabilities would be time-independent variables that specify the fraction of each sub-population of the ensemble; this approach is called a mixture model. Marginalization would then involve summing the likelihoods for the different sub-populations, weighted by the probabilities of those subpopulations. Consequently, during the inference procedure, the probabilities of the sub-population occupancies would then become model parameters that are also inferred using Bayes’ rule.

Additionally, it is possible to have a time-dependent system with hierarchical transitions between the different sub-populations. In this case, the probabilities of each 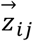 in the 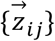 would not be constant for each sub-population, as they would be for a mixture model, but would instead depend upon the sub-population of the previous data point 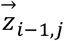 and a K × K transition matrix, 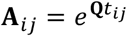, where **Q** is the rate matrix that depends upon the set of rate constants for transitioning between the K different states, and *t*_*ij*_ is the time that has elapsed since the previous data point, which may not necessarily be equal to *τ* (Fig. S1B). Here, marginalization is efficiently performed with the forward-backward algorithm (49) and the steady-state probabilities, as calculated from the rate constants for the kinetic scheme under consideration, for instance by using the diagram method (50), are used to set the initial probability of each 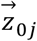. In total, this approach amounts to a hierarchical, continuous-time, ensemble HMM for sub-temporal resolution systems, where inference is performed directly upon the rate constants, instead of the transition probabilities. Consequently, this approach can handle shuttering of the laser light source in fluorescence microscopy experiments or other types of irregular spacing of data points, sub-temporal resolution data, and population-level analyses with non-parametric posterior distributions, which can be used to ascertain the underlying thermodynamic landscape of the mesoscopic ensemble.

To highlight this hierarchical approach, consider a single-molecule fluorescence microscopy experiment in which a fluorophore-labeled biomolecule transitions between two states, 1 and 2, with forward and reverse rate constants *k*_12_ and *k*_21_, respectively (Fig. 4A). Such fluorescence microscopy experiments often suffer from photophysical phenomena such as fluorophore photoblinking, in which a fluorophore temporarily transitions into a long-lived, ‘dark’, excited molecular electronic state and thus transiently stops fluorescing, or fluorophore photobleaching, in which a fluorophore that has transitioned into an excited molecular electronic state undergoes a photochemical reaction and permanently stops fluorescing (51). Often, the transition rates into and out of the dark states responsible for photoblinking are faster than the time resolution of techniques such as wide-field, fluorescence microscopy. As a result, instead of detecting a steady level of fluorescence intensity from the fluorophore, sub-temporal-resolution transitions between fluorescent and dark states of the fluorophore manifests as an extra, and often dominant, source of ‘noise’ in the single-molecule fluorescence intensity signal trajectory (Fig. 4B, inset). Intense experimental effort has gone into minimizing these photophysical effects, including the use of fluorophores, such as Cy3B, that have been chemically altered so as to minimize transitions to dark states (52); elaborate excitation laser modulation schemes, such as triplet-state relaxation (T-Rex) and dark-state relaxation (D-Rex) schemes, that minimize transitions to higher-order dark states (53, 54); photostabilizing additives, such as Trolox, that accelerate transitions out of dark states (55, 56); and fluorophore-photostabilizer conjugates, such as Cy3-and Cy5-triplet state quencher conjugates, that accelerate transitions out of dark states (57, 58). Here, we show how extending BIASD with the hierarchical HMM described above allows us to computationally overcome these photophysical effects.

**Figure 4:**
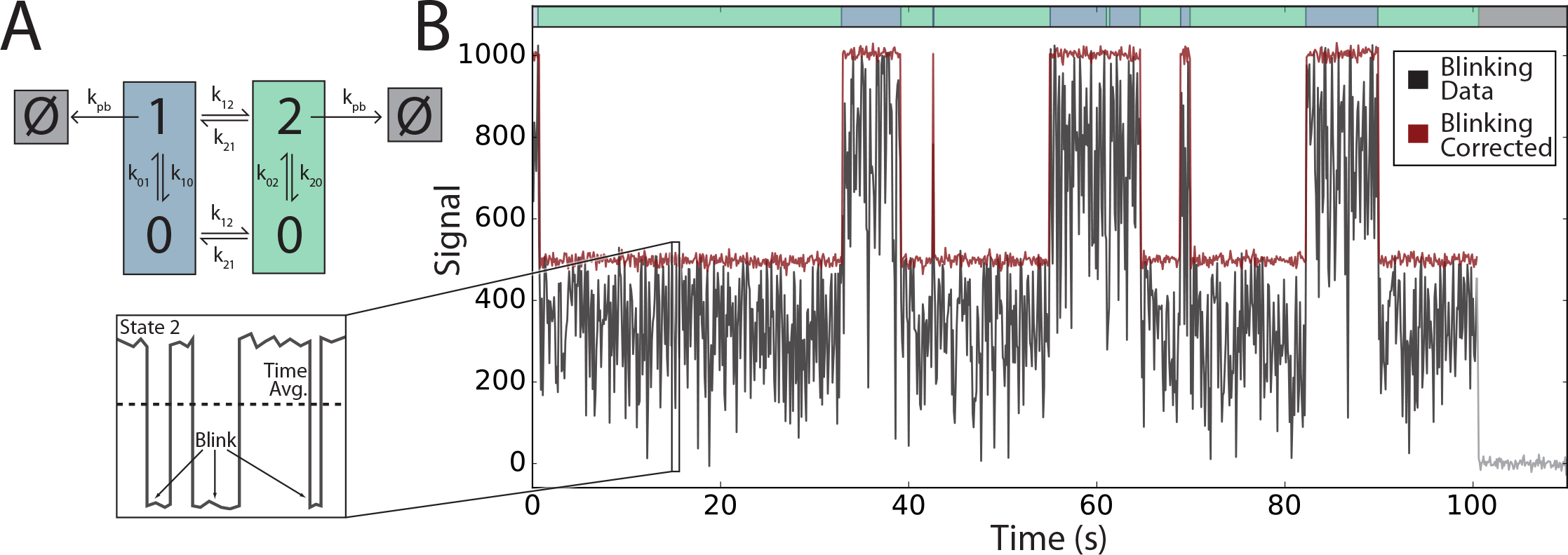
(A) Kinetic mechanism used to simulate an experimental system in which a biomolecule transitions between two conformational states that can each transition, with sub-temporal resolution dynamics, into and out of a photoblinked state, until eventually photobleaching. Exact values of the rate constants used in the simulation are given in the *Supporting Information.* (B) Plot of simulated signal trajectory and an estimated, corrected fluorescence intensity signal trajectory in the absence of photoblinking. The green-and blue-colored regions at the top of the plot denote the time spent in states 1 (blue) and 2 (green). The signal corrupted by sub-temporal resolution photoblinking that was analyzed by BIASD is shown in black; the inset shows a cartoon of a single *τ* where fast transitions are taking place between the fluorescent and photoblinked state 2. After analysis with BIASD, the MAP solution of the posterior probability distribution was used to generate a Viterbi-idealized path, which is plotted in red with noise added back from the MAP value of σ; this is a plausible corrected fluorescence intensity signal trajectory in the absence of photoblinking. While one excursion to state 2 is not present in this Viterbi path, the forward-backward algorithm used in the HMM analysis takes it into consideration.

To demonstrate this ability, we simulated the kinetic scheme shown in Fig. 4A, where a fluorophore-labeled biomolecule transitions between conformational states 1 and 2 with signal values of ∊_1_ and ∊_2_, respectively. However, in this simulation, both of these states can rapidly transition into and out of a photoblinked state, denoted 0 with signal value ∊_0_ = 0, at rates much faster than the time resolution of the simulated data. These dynamics continue until the system eventually transitions into a photobleached state, denoted Ø with signal value ∊_ø_ = 0. A similar situation has been recently investigated by Chung *et al.* to analyze FRET photon trajectories reporting on the sub-temporal resolution folding and unfolding dynamics and photoblinking dynamics of the villin subdomain protein (20). As expected, by analyzing this simulation using this hierarchical approach, the posterior probability distribution of the parameters describing the fluorescence emission from each sub-population (∊_1_, ∊_2_, *σ*, *k*_0_, *k*_01_, *k*_20_, and *k*_02_; see Fig. S1B), as well as the rate constants describing the transitions between states 1 and 2 (*k*_12_, and *k*_21_; see Fig. 4A), were all found both accurately and precisely as the parameter values used for the simulation fall within the inferred 95% credible intervals (Fig. S3). To provide visual intuition into this result, we also have shown the Viterbi-idealized path from the maximum *a posteriori* (MAP) estimate of the model parameters in order to show the most likely fluorescence intensity signal trajectory in the absence of photoblinking (Fig. 4B). Detection noise from the marginalized posterior distribution of σ was added to this path to show what the data might have looked like in the absence of photoblinking. Regardless, we note that this particular path is essentially a point estimate of the 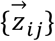, whereas, by marginalizing out all of the 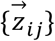 during the inference procedure, we have actually considered all the other possible paths, regardless if a transition is missed in the Viterbi path. As such, the posterior probability distribution of the model parameters is a more encompassing result (Fig. S3). Finally, we note that the hierarchical HMM treatment that we present here is general and applicable to not just two, but to any number of K sub-populations.

### Conclusion

By analyzing the fraction of time that a single-molecule spends in each state of a defined kinetic scheme during each *τ* in a signal trajectory, BIASD adopts a fundamentally different approach to the analysis of time-resolved, single-molecule experiments than that which has been traditionally employed by methods that idealize the trajectories (*e.g.*, signal thresholding, HMMs, *etc.*). Using computer-simulated and experimentally observed data, we have demonstrated that this powerful approach enables BIASD to accurately and precisely infer the rate constants of a two-state kinetic scheme as well as the signal values corresponding to these two states, even when the rates of transitions between the states are orders of magnitude larger than the time resolution of the signal trajectories. When used to analyze experimental E_FRET_ trajectories reporting on the dynamics of single PRE^−A^ complexes recorded as a function of temperature (9), BIASD allowed us to infer the thermodynamic activation parameters characterizing the transition-state energy barriers along the GS1→GS2 and GS2→GS1 reaction coordinates, which had thus far remained inaccessible to traditional smFRET data analysis approaches. Moreover, we have demonstrated that a straightforward extension of the BIASD framework enables the kinetics of experimental systems exhibiting multiple sub-populations of molecular properties to be accurately and precisely inferred.

It is important to note that the BIASD framework is general and can be applied to any experimentally observed signal trajectory that exhibits stochastic transitions between distinct states, regardless of the nature or the origin of the signal. Thus, BIASD can be used to temporally resolve data collected using virtually any time-resolved single-molecule experimental method, including single-molecule fluorescence microscopy, force spectroscopy, conductance, and tethered particle motion methods. Moreover, although here we have developed BIASD to analyze single-molecule signal trajectories, we have not consider the temporal ordering of the data. Consequently, in addition to analyzing individual single-molecule signal trajectories, BIASD can also be used to analyze the distribution of fractional occupancies observed across an entire ensemble of individual molecules during a given *τ*. This could allow non-equilibrium phenomenon to be monitored across an ensemble of single molecules, time period by time period (*e.g.*, stopped-flow delivery of a ligand, substrate, cofactor, or inhibitor to an enzyme or other biomolecule). In addition, BIASD can be expanded to include the time evolution of the state occupation probabilities (*c.f*, Eqn. 2 of the *Supporting Information*), or to incorporate time dependence into the model parameters {*k*}, {∊}, and {σ} (*e.g.*, the varying of {∊} in single-molecule particle tracking experiments).

Regarding the performance of BIASD on experimental data, we note that the rate constants and signal values of a system can be more precisely inferred from experiments that collect higher SNR data, because then there is less uncertainty in the time-averaged fractional occupancies of the signal trajectories. Therefore, somewhat counterintuitively, sub-temporal-resolution dynamics can to some degree be more precisely inferred from signal trajectories recorded with lower time resolutions but higher SNRs (*e.g.*, due to better photon conversion efficiencies on an electron-multiplying charge-coupled device), than those recorded with higher time resolutions but lower SNRs. Additionally, although we have focused the current work on the most widely applicable case of a Markovian, two-state system in which the noise of the signal can be modeled using a normal distribution, the Bayesian inference-based framework underlying BIASD can be readily extended to non-Markovian dynamics (21, 59), N-state kinetic schemes (60, 61), or systems in which the noise of the signal can be modeled using distributions other than a Normal distribution (18, 62). It should be noted, however, that such developments will come with added computational expenses. To facilitate the analysis of single-molecule data using BIASD, as well as to enable the future extension of BIASD along the lines described here, we have made the BIASD source code available at http://github.com/ckinzthompson/biasd. The source code is written in Python and integrated with computationally intensive functions provided in C as well as in CUDA (for GPU-based computation) in order to balance accessibility with high-performance.

## Methods

### Simulating Signal Trajectories

State trajectories were simulated with the stochastic simulation algorithm (36); briefly, sequential random lifetimes were drawn from exponential distributions with the specified rate constants, and subsequent states were chosen randomly according to the splitting probabilities. A random starting point for the initiation of the trajectory (t = 0 sec) was selected with a uniform distribution from the first lifetime. The fractional occupancies of each state during each sequential *τ* were then calculated from the sequence of lifetimes. The resulting fractional occupation *versus* time trajectories were turned into signal trajectories by computing *μ*, and then adding normally distributed noise with standard deviation, σ. Simulations of the titration experiment were performed such that ∊_1_ = 0, ∊_2_ = 1, σ_1_ = σ_2_ = 0.04, 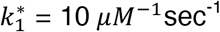, *k*_2_ = 10 sec^−1^, and *τ* = 0.1 sec, and each signal trajectory was 600 data points in length. Parameters for the simulations with the transitioning subpopulations are given in the *Supporting Information*.

### Bayesian Thresholding Analysis

Signal trajectories were idealized by thresholding any measurement period with signal less than (∊_1_ + ∊_2_)/2 = 0.5 into state 1, and otherwise into state 2. Rate constants from the i^th^ state to j^th^ state were then calculated as *k*_*ij*_ = −ln (1 − *p*_*ij*_)/*τ* where *p*_*ij*_ is the transition matrix from the idealized trajectory (2). Credible intervals for the transition probabilities, and rate constants were calculated with uniform prior distributions (2). The joint posterior probability distributions of ∊_1_ and 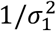, and ∊_2_ and 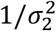 were inferred using the analytical formulas for Bayesian inference with a joint normal-gamma prior probability distribution using those data points that were idealized into the respective states (6). The marginalized posterior distributions of the ∊_*i*_ and 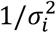 (T and gamma distributions, respectively) were used to calculate means and 95% credible intervals for each parameter (6). An aggregate σ was then calculated by weighting 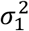 and 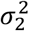 by the fraction of data points idealized into each state and taking the square root of their sum.

### Maximum Likelihood HMM Analysis

Signal trajectories were analyzed using a two-state, discrete, maximum likelihood HMM with normal distribution emissions using the expectation maximization, and forward-backward algorithms (6, 49). Each trajectory was analyzed with 20 randomized restarts, including one initialized at the simulated values, until the likelihood of each restart converged to a relative value of 10^−10^. From these, the point-estimate with the greatest likelihood was used in subsequent analyses. Rate constants were calculated directly from the transition probability matrix point-estimate, and an aggregate standard deviation was calculated as described above for Bayesian thresholding.

### BIASD Analysis

Adaptive, Gauss-Kronrod (G10, K21) quadrature was used to numerically integrate the BIASD likelihood function on an Nvidia GeForce 1080 GTX graphics card; the likelihood of each data point took about 1 μsec to compute. The posterior probability distribution was sampled using *emcee,* an ensemble, affine-invariant MCMC method (34, 35). For each trajectory, 100 MCMC walkers were employed to draw 2000 samples each, and the first 1000 samples were discarded to burn in the chain. From the remaining samples, independent samples were chosen spaced apart by the maximum parameter autocorrelation time, and credible intervals and means were calculated from these samples. It took approximately two minutes to sample a single, 600-data-point signal trajectory, such as those used in the computer simulated titration.

### PRE^−A^ E^FRET^ Analysis

Previously published Cy3 and Cy5 fluorescence intensity, I_Cy3_ and I_Cy5_, *versus* time trajectories from the PRE^-A^ complex from the study by Wang and coworkers (9) were transformed into E_FRET_ trajectories by calculating E_FRET_ = I_Cy5_/(I_Cy3_ + I_Cy5_) at each measurement period. Outliers where E_FRET_ < −0.4 or E_FRET_ > 1.4 were clipped. The number of E_FRET_ trajectories in the 22, 25, 28, 31, 34, and 37 °C datasets were 490, 456, 435, 452, 270, and 459, respectively. Uniform distributions were used for the prior probability distributions. The first and second moments, *E*[*k*] and *E*[*k*^2^], of the marginalized posterior probability distributions for *k*_*GS*1_ or *k*_*GS*2_ were used to infer the values of Δ*H*^‡^, Δ*S*^‡^, and a precision *λ* using Bayesian inference with the likelihood function 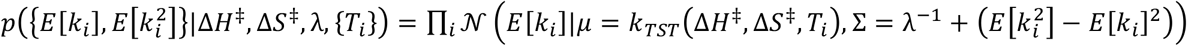, where *N* is the normal distribution with mean *μ* and variance ∑, where *k*_*TST*_(Δ*H*^‡^,Δ*S*^‡^,*T*_*i*_) is the rate constant calculated at temperature *T*_*i*_ with transition state theory, and where *i* indexes the set of temperatures. The resulting posterior probability distribution for Δ*H*^‡^, Δ*S*^‡^, and *λ* was sampled using MCMC from which credible intervals and means were calculated.

## Acknowledgements

The authors would like to thank Prof. Jan-Willem van de Meent for his comments on an early version of this manuscript. This work was supported by two NIH-NIGMS grants (R01 GM084288 and R01 GM 119386), an American Cancer Society Research Scholar Grant (RSG GMC-117152), and a Camille Dreyfus Teacher-Scholar Award (DRFSCH CU11-0665) to R.L.G. C.D.K. was supported by the Department of Energy Office of Science Graduate Fellowship Program (DOE SCGF), made possible in part by the American Recovery and Reinvestment Act of 2009, administered by ORISE-ORAU under contract number DE-AC05-06OR23100, and by Columbia University's NIH Training Program in Molecular Biophysics (T32-GM008281).

## Author Contributions

C.D.K. conceived of and performed the research; C.D.K. and R.L.G. discussed results and wrote the manuscript.

